# microRNA-451a regulates colorectal cancer radiosensitivity

**DOI:** 10.1101/136234

**Authors:** Rebecca Ruhl, Shushan Rana, Katherine Kelley, Cristina Espinosa-Diez, Clayton Hudson, Christian Lanciault, Charles R Thomas, Liana V Tsikitis, Sudarshan Anand

## Abstract

Colorectal cancer (CRC) is a leading cause of cancer-related death. The responses of CRC to standard of care adjuvant therapies such as radiation or chemotherapy are poorly understood. MicroRNAs (miRs) are small non-coding RNAs that affect gene expression programs in cells by downregulating specific mRNAs. In this study, we discovered a set of microRNAs upregulated rapidly in response to a single 2 Gy dose fraction of γ-radiation in a mouse colorectal carcinoma xenograft model. The most upregulated candidate in our signature, miR-451a inhibits tumor cell proliferation and attenuated surviving fraction in longer-term cultures. Conversely, inhibition of miR-451a increased proliferation, tumorsphere formation and surviving fraction of tumor cells. Using a bioinformatics approach, we identified four genes-CAB39, EMSY, MEX3C and EREG as targets of miR-451a. Transfection of miR-451a decreased both mRNA and protein levels of these targets. Importantly, we found miR-451a expression was decreased with tumor stage in a small subset of CRC patients. Finally, analysis of a TCGA colorectal cancer dataset reveals that the CAB39 and EMSY are upregulated at the protein level in a significant number of CRC patients and correlates with poorer overall survival. Taken together, our data indicates miR-451a influences the radiation sensitivity of colorectal carcinomas.

## Introduction

In 2016, an estimated 134,000 patients will be diagnosed with colorectal cancer. Among the rectal cancer subset, patients with locally advanced disease, staged as T3-T4 and or node positive, receive neoadjuvant chemoradiation therapy (CRT) and subsequent surgery ^1,2^. The standard of care is still surgical resection with total mesorectal excision that may result to significant quality of life issues ^3^. Many patients even after neoadjuvant chemoradiation may never be rendered resectable. Response to CRT is an independent predictor of overall survival in colorectal cancer ^4^ highlighting the importance of improving CRT response rates. It is known that several tumor intrinsic factors govern responses to CRT including specific gene expression programs with distinct significance ascribed to microRNAs (miRs) ^5,6^. miR-processing machinery is frequently mutated in colorectal cancers (TCGA, 2016 provisional), and miRs have been implicated in several pathological processes associated with colorectal cancer progression including cancer stemness and epithelial-to-mesenchymal transition (EMT) ^7,8^. Emerging evidence suggests that microRNAs (miRs) modulate gene expression programs in response to radiation and confer variable sensitivity and efficacy to modern high dose ionizing radiation therapy ^9,10^. In this context, we have identified miR-451a as a tumor suppressor miRNA in colorectal adenocarcinoma whose presence correlates with increased radiation efficacy. Through gain and loss-of-function studies, we show that miR-451a is a negative regulator of proliferation in CRC and likely mediates its effects by targeting CAB39 and EMSY. We believe our work highlights the potential for using miRs and its target genes in predicting radiation responsiveness of CRC while also illustrating potential avenues for restoring radiation sensitivity in poorly responding tumors.

## Results

### Early and late radiation responsive miRs in CRC

To identify miRs that are regulated by radiation in CRC, we implanted either HCT-116 or SW-480 xenografts into nude mice. After the tumors reached a 300mm^3^ volume, we treated the mice with a single 2 Gy focal radiation. Tumors were harvested at either 6 h or 48 h post-RT and RNA was extracted to generate the initial *in vivo* miR profile using Affymetrix microRNA arrays. Using a 2-fold regulation in both cell lines as a cut-off, we identified a group of two miRs that were upregulated and two miRs that were downregulated at 6h (Fig 1a). The top candidate in this profile, miR-451a was validated using qRT-PCR across three different human CRC cell lines (Fig 1b) grown as subcutaneous xenografts.

**Figure 1.**
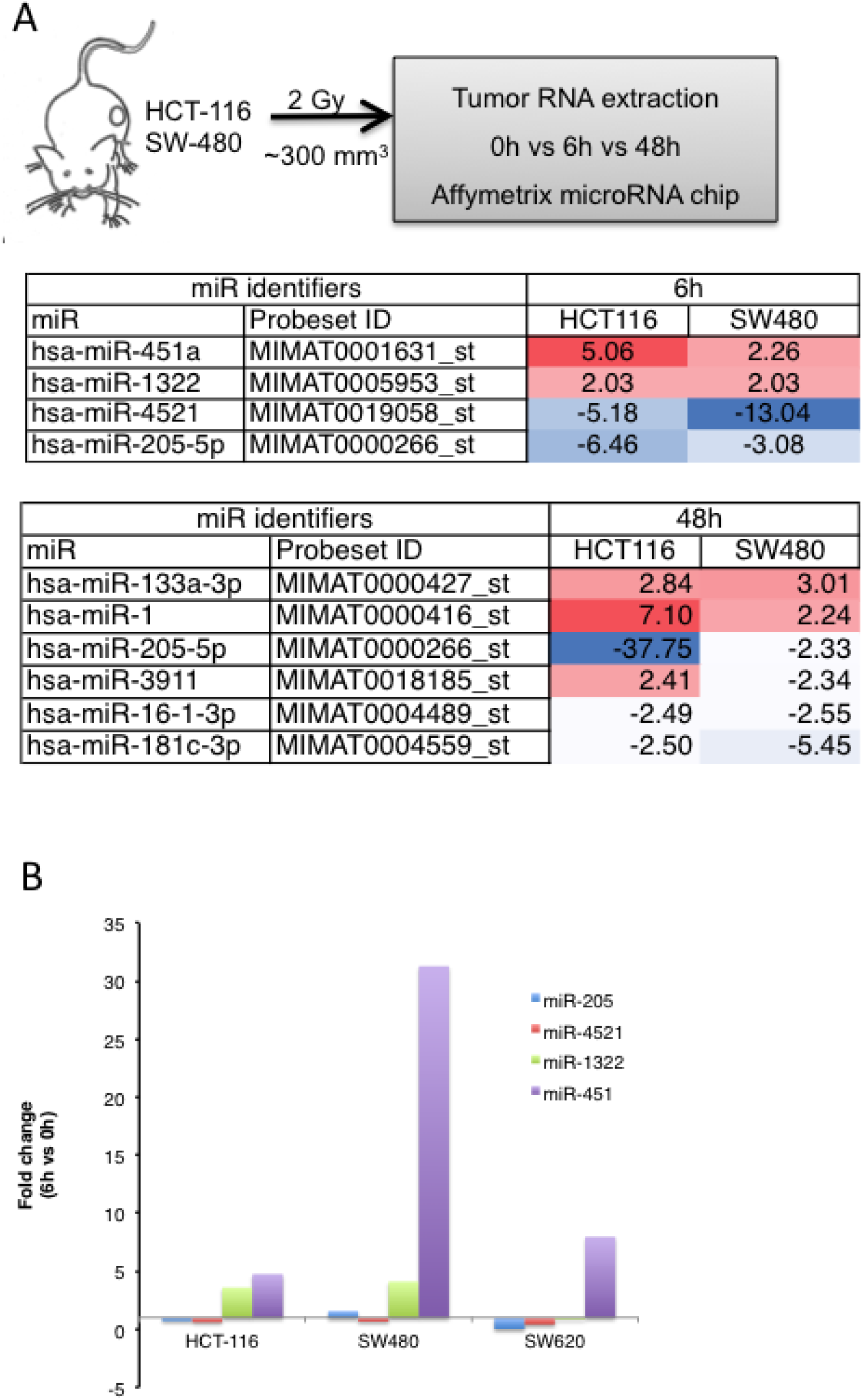
miR-451a is robustly induced by radiation in human colorectal carcinoma cells in vitro and in vivo. A) Design of the screen for miRs induced by radiation. Heatmap depicts miRs with more than 2 fold change from the affymetrix microRNA array v4.0 across both tumor types. B) Levels of the 2 most upregulated and downregulated miRs 6h post 2 Gy radiation were validated by qRT-PCR using specific Taqman probes for each microRNA. Mean fold change after normalization to a housekeeping RNA, RNU48, is depicted.

### miR-451a functions as a tumor suppressor miR in colorectal cancer

To understand the function of miR-451a, we first performed gain of function studies in vitro with HCT-116 cells. Ectopic expression of miR-451a decreased proliferation in 2D culture (Fig 2a), tumorsphere assay (Fig 2b) and in surviving fraction colony formation assay (Fig 2c). We noted that while miR-451a alone had a significant effect on the phenotypes, there was also slight additive effect with radiation, especially at the 5 Gy dose (Fig 2c). This inhibition of proliferation and clonogenic survival was also confirmed in CT-26 cells, a mouse colorectal adenocarcinoma cell line (Supplementary Figure 1). Conversely, inhibition of miR-451a enhanced proliferation in 2D (Fig 3a) and tumorsphere assays (Fig 3b) almost negating the effects of a 2 Gy dose of radiation. The preservation of survival was most robust in the 2 Gy population with a log fold increase in clonogenic survival, but was not evident at higher dose radiation (Fig 3c). Similar to these findings, miR-451a transfection also enhanced the responses to 5-FU a commonly used chemosensitizer in colorectal cancer (Supplementary Fig 2). Taken together, these studies indicate that miR-451a regulates proliferation of colorectal cancer cells.

**Figure 2.**
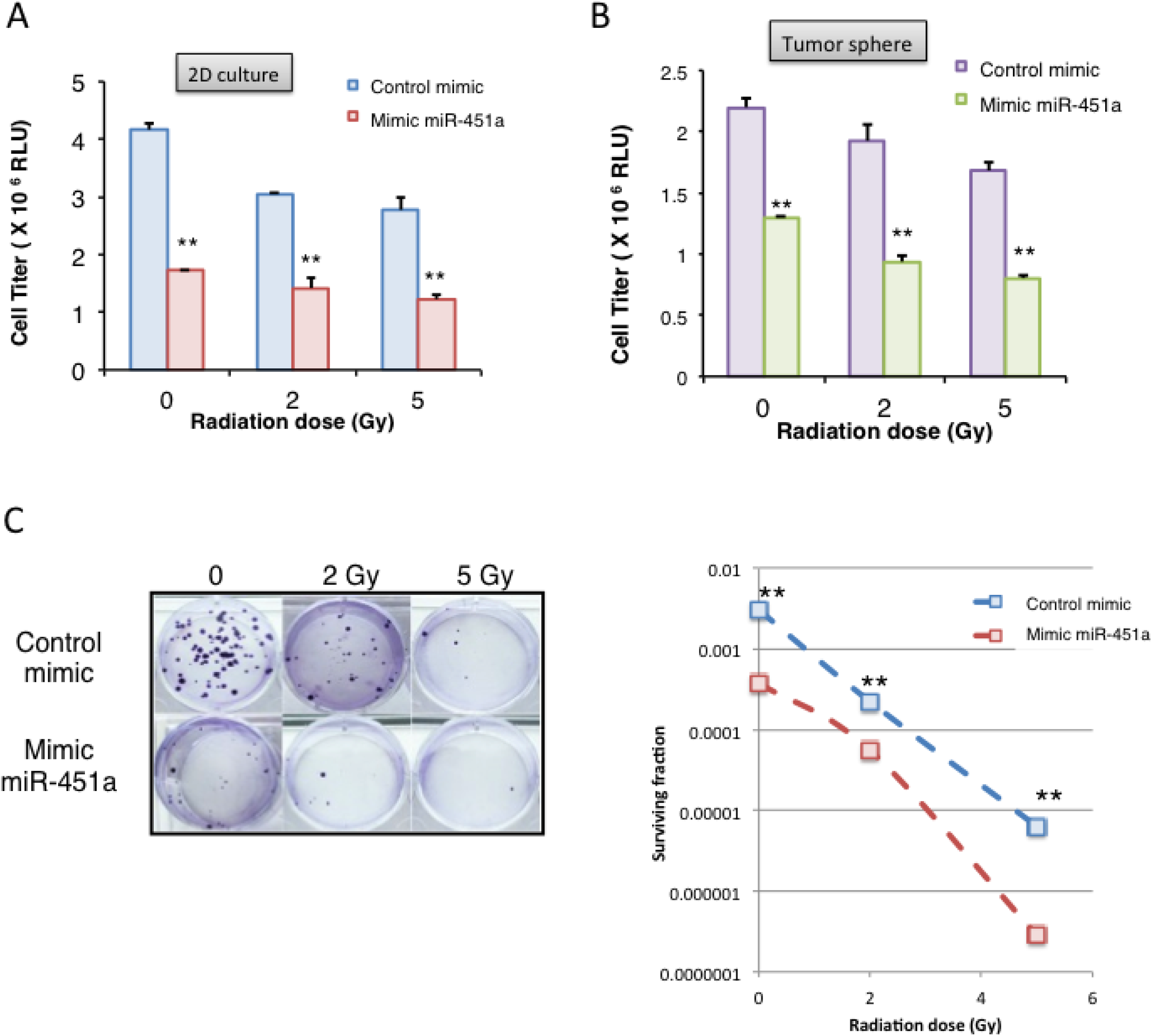
Ectopic expression of miR-451a inhibits proliferation and clonogenic survival of HCT-116 cells. HCT116 were transfected with a miR-451a mimic or a control mimic. Proliferation was analyzed 48 hours after radiation in A) 2D and B) 3D cultures with the indicated doses. Bars depict mean ± s.e.m. of triplicate wells. ** indicates P< 0.01 on a twotailed Student’s T-test. C) 12-14 days after plating, cells were fixed and stained with crystal violet and colonies were counted. Surviving fraction was calculated based on the colony numbers normalized to the plating efficiency. Mean of triplicate wells is plotted. * indicates P< 0.05 and ** indicates P< 0.01on a two-tailed Student’s T-test

**Figure 3.**
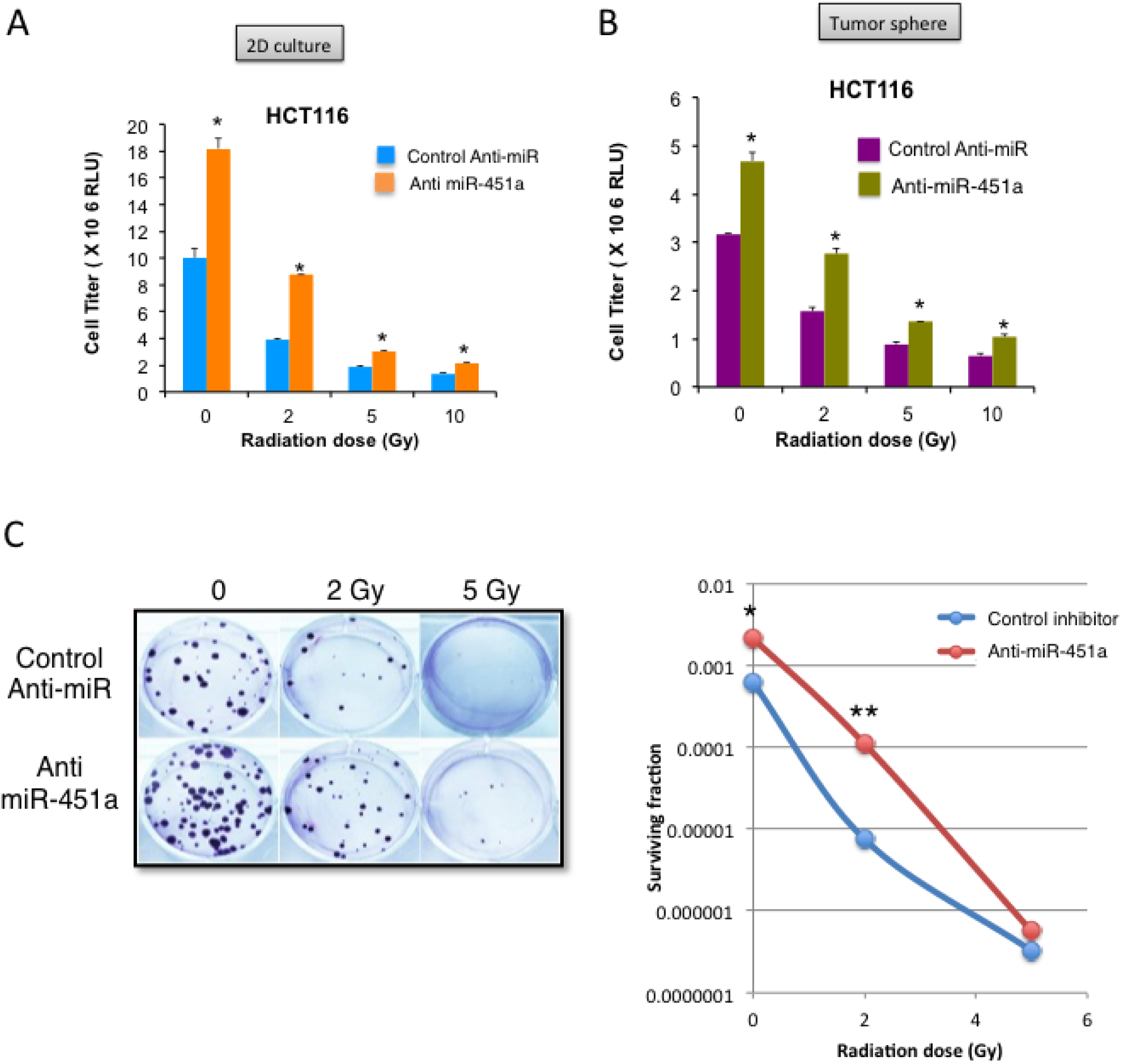
miR-451a inhibition increases tumor cell proliferation and clonogenic survival. HCT116 were transfected with a miR-451a inhibitor (anti-miR-451a) or a control anti-miR. Proliferation was analyzed 48 hours after radiation in A) 2D and B) 3D cultures with the indicated doses. Bars depict mean ± s.e.m. of triplicate wells. ** indicates P< 0.01 on a twotailed Student’s T-test. C) 12-14 days after plating, cells were fixed and stained with crystal violet and colonies were counted. Surviving fraction was calculated based on the colony numbers normalized to the plating efficiency. Mean of triplicate wells is plotted. * indicates P< 0.05 and ** indicates P< 0.01on a two-tailed Student’s T-test

### miR-451a targets genes involved in metabolism and DNA repair pathways

To elucidate the relevant miR-451a targets in relation to radiation and colorectal cancer, we used the miRwalk algorithm to combine data from multiple target prediction databases and identified 14 genes as putative targets of miR-451a. We further narrowed this list by filtering genes with a known role in human colorectal cancer and/or cellular response to ionizing radiation that resulted in a group of four genes -CAB39, EMSY, EREG, and MEX3C (Supplementary Figure 3). We used a miR-Trap assay to pull down mRNAs enriched at the RNA induced silencing complex in cells transfected with miR-451a and validated these four target mRNAs were enriched at the RISC (Fig 4a). Indeed transfection of HCT-116 cells resulted in significant downregulation of all four target genes at the mRNA level (Fig 4b). We validated the decrease in mRNA levels resulted in a decrease in protein expression for CAB39 and EMSY (Fig 4c and Supplementary Fig 4). Moreover, siRNA mediated silencing of two of the four targets EMSY and CAB39 in HCT-116 cells recapitulated the miR-451a induced inhibition of proliferation (Fig 4d). These experiments suggest that miR-451 mediated decrease of EMSY and CAB39 is the likely reason for the decrease in proliferation.

**Figure 4.**
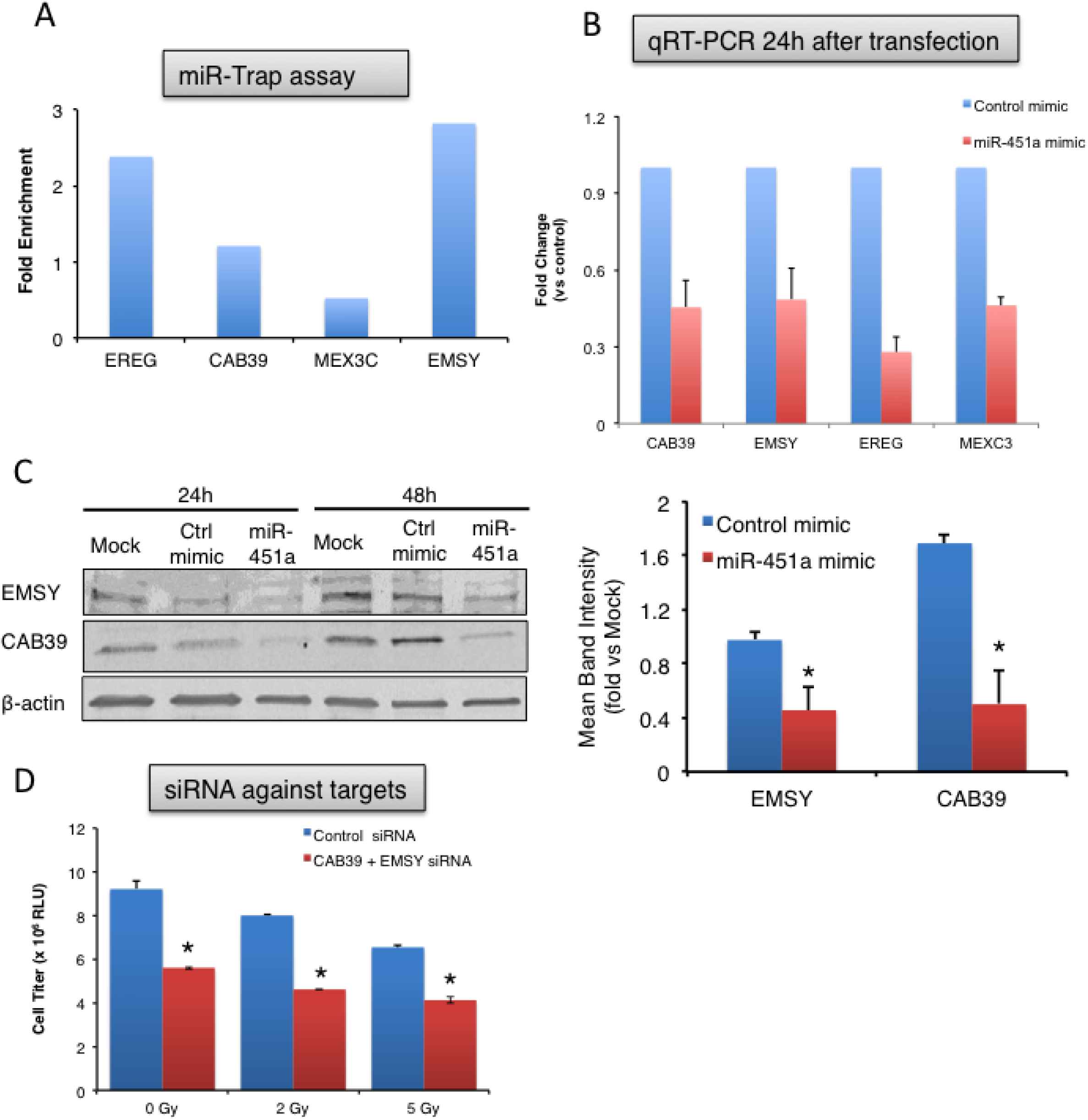
miR-451a targets genes involved in cell cycle and cellular stress responses. A) miR-TRAP assay depicting enrichment of target mRNAs immunoprecipitated from HCT116 cells co-transfected with a mutant RISC complex plasmid and a miR-451a mimic or a control miR mimic. Fold enrichment over pre-IP mRNAs is depicted. One of two independent experiments. B) qRT-PCR of the miR-451 targets in HCT-116 cells at 24h after transfection. C) Western blot for EMSY and CAB39 in HCT-116 cells at 24h and 48h after transfection of miR-451a compared to control mimic or mock transfection. Right panels show quantitation of two independent experiments. Full blot is shown in Supplementary Figure 4 D) siRNA mediated silencing of EMSY and CAB39 phenocopies the miR-451a effects in HC-116 proliferation. * P<0.05, Student’s T-test.

### miR-451a and target expression in colorectal cancer patients

To assess if miR-451a was relevant in human colorectal cancer, we measured the miR levels in a colorectal carcinoma tissue array (Supplementary Fig 5 and Fig 5a). While we did not see a statistically significant difference, we observed a trend towards a decrease in miR levels with an increase in tumor stage. Moreover, analysis of the TCGA database (Provisional colorectal carcinoma, n=639 patients) showed that CAB39 and EMSY protein levels were found to be upregulated in 14% and 6% of cases, respectively (Fig 5b). Interestingly, upregulated expression of these genes conferred a significantly reduced 3 year overall survival (Fig 5c).

**Figure 5.**
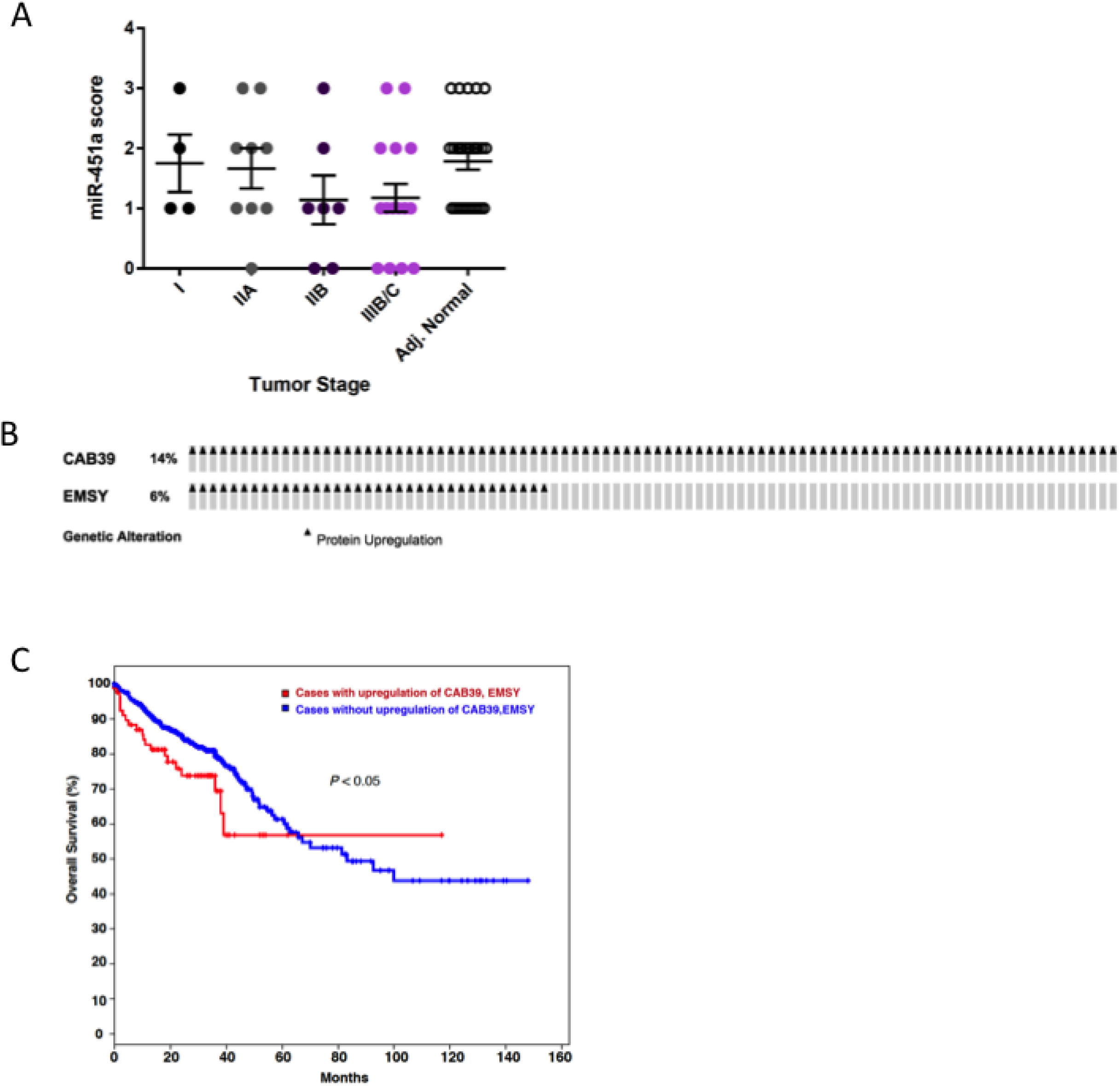
Regulation of miR-451a and target genes in human colorectal cancer. A) miR-451a levels as measured by in situ hybridization across different stages of a human colorectal carcinoma tissue array. Error bars show SEM. B) CAB39 and EMSY protein levels upregulated in a subset of patients in the TCGA Colorectal Adenocarcinoma (provisional) cohort of 633 samples of which 498 were assayed for protein expression levels. C) Kaplan Meier plot showing survival of cases with upregulation of CAB39, EMSY compared to the cases without upregulation. P value from a log-rank test.

## Discussion

Several recent studies have shown the utility of miRs in the diagnosis and classification of CRC^7,8,11-14^. There are ongoing prospective clinical trials evaluating miR based classifiers such as a 24-miR signature in lung cancer diagnosis ^15^ (Gensignia) and a miR signature in prostate cancer screening ^16^ (Exiqon). These trials highlight the feasibility and translational potential of miR based classifiers. We identified a group of miRs that are responsive to radiation in mouse xenograft models and focused on characterizing miR-451a as one of the robust early response miRs in CRC. Our data suggests that miR-451a behaves as a tumor suppressor in CRC cell lines in vitro. We also identify putative targets of miR-451a and show that expression of two of the targets correlate with overall survival in human colorectal cancer. Taken together, our observations suggest that miR-451a modulation of CAB39 and EMSY target genes could alter radiation sensitivity of human CRC.

miR-451a has been found to inhibit cell proliferation and drug responses in other malignancies. For example, miR-451a was found to affect proliferation and sensitivity to tamoxifen in breast cancer via targeting of the macrophage migration inhibitory factor (MIF) ^17^. Similarly, miR-451a was shown to be tumor suppressive in gastric cancer by affecting the PI3K/mTOR pathway ^18^. In other tumor types, it has been shown that miR-451a expression is downregulated in the tumor cells in a manner consistent with a tumor suppressor function ^19,20^. Interestingly, miR-451a appears to increase radiation responses in nasopharyngeal carcinoma cells ^21^ and lung adenocarcinoma cells ^22^. Our observation in CRC cell lines suggests a function similar to the tumor suppressive role that have been documented in other cancers by these studies.

Using a bioinformatics approach, we narrowed down the targets of miR-451a to a group of 14 genes which we manually narrowed down to 4 genes –CAB39, EMSY, EREG and MEX3C based on either a known role in colorectal cancer or radiation responsiveness in other cancer types. Calcium binding protein (CAB) 39 has been previously shown to be a target of miR-451a in human glioma ^23^. This protein is thought to affect STK11 activity and localization thereby influencing the PI3K/AKT signaling pathway. EMSY is a transcriptional repressor that associates with BRCA2 and is often amplified in breast and ovarian cancers ^24^. Functionally, EMSY colocalizes and forms foci with histone ƔH2AX in response irradiation. Importantly, breast cancer patients with EMSY amplifications have poorer overall survival. Taken together, these functions suggest that modulation of EMSY by miR-451a may have significant impact on radiation responses and tumor cell survival. Indeed, consistent with the breast cancer dataset, our analysis of the TCGA colorectal cancer dataset (Fig 5b-c) indicates that EMSY as well as CAB39 are increased at the protein level in a subset of patients and associated with worse outcome. Epiregulin (EREG) is a known ligand of the EGF family and regulates several key processes including cellular proliferation, inflammation, angiogenesis and wound healing ^25^. While it has been proposed as a biomarker for monitoring responses to cetuximab in colorectal cancer ^26^, it is unclear whether there is a function for EREG specifically in the context of radiation responses of these tumors. MEX3C has been identified as a ubiquitin ligase as well as an RNA binding protein that modulates the levels of HLA-A allotypes ^27^. Interestingly, it is part of a group of genes that suppress chromosomal instability in colorectal cancer ^28^ thereby likely contributing to tumor drug resistance. Our data suggests that miR-451a mRNA binds to all four of these mRNAs (Fig 4a) and downregulates their expression at the RNA and protein levels. Given the low expression levels of the EREG and MEX3C in patient samples, we chose to focus on CAB39 and EMSY and found that there was a difference in overall survival in the TCGA in patients with upregulation of CAB39 and EMSY. We also noticed a trend towards decreased miR expression in patients with more advanced disease compared to adjacent normal tissue. We must emphasize that our small patient numbers preclude drawing stronger conclusions regarding the miR-451a and target levels in human disease but rather lead us to hypothesize that tumors with increased miR-451a levels will respond better to CRT and result in enhanced overall survival.

Our work shows that miR-451a is a radiation-induced miR in CRC and identifies novel targets of miR-451a that may contribute to radiation responses. Further work is required to validate these observations in larger numbers of patients as well as elucidate the mechanistic basis of miR-451a mediated decrease in proliferation via these target pathways. We envision that our data herein will enable the development and validation of miR and/or target biomarkers that predict radiation responsiveness as well as provide strategies for enhancing the effectiveness of chemoradiation in colorectal cancers.

## Methods

### RNA extraction, RT-PCR, miR Profiling

RNA was extracted using miRVana microRNA isolation kit (Ambion/Life Technologies). Affymetrix microRNA array v4.0 profiles were generated by the OHSU Genome Profiling Core facility. Individual RT-PCRs were performed using predesigned TaqMan Assays for mature miRs, primary miRs or mRNAs (Applied Biosystems) on a Vii-7 real time PCR platform (Applied Biosystems) according to manufacturer’s instructions. Data was normalized to internal control small RNA RNU48 or U6 small RNAs. mRNAs were normalized to either β-actin or GAPDH.

#### Cell Culture and Reagents

HCT-116 cells (ATCC) were cultured in McCoy’s supplemented with 10% FBS and antibiotics. CT-26 cells (ATCC) were cultured in RPMI-1640 medium supplemented with 10% FBS and antibiotics. SW480 and SW620 cells (ATCC) were cultured in Leibovitz’s L-15 Medium with 10% FBS, antibiotics under 0% CO_2_ conditions. Cells were tested and found negative for mycoplasma contamination before use in the assays described.

#### Transfections

Cells were reverse transfected with miR-451a-5p mimics, inhibitors, selected siRNAs against CAB39 and EMSY and their respective controls purchased from Life Technologies using Lipofectamine RNAiMAX (Invitrogen) according to manufacturer’s instructions.

#### Colony Formation Assay

Cells were transfected with miR-451a-5p mimic or control mimic for 16 hours. Then cells were plated (100 or 200 cells for 0Gy, 200 or 400 cells/well for 2GY and 400 or 800 cells per well for 5Gy) in triplicate in a 6-well plate. Cells were irradiated 4 hours after plating, 0Gy, 2Gy or 5Gy. Two weeks after plating, cells were fixed and stained with crystal violet and colonies were counted. Surviving fraction was calculated based on the colony numbers normalized to the plating efficiency.

#### In vivo assays

All animal work was approved by the OHSU Institutional Animal Use and Care Committee. Animal experiments were performed in accordance with the OHSU IACUC guidelines and regulations. Immune compromised 8-10 week old nu/nu mice purchased from Jackson Labs were injected subcutaneously with 1 million mycoplasma-negative tumor cells in Matrigel (BD) per flank. Tumor growth was measured with calipers, with volume computed as ½ * Length * Width^2^. Mice were randomized into groups once the average tumor volume reached 80mm^3^, approximately 6 days after injection.

#### Irradiation

Cells or mice were irradiated on a Shepherd^137^ cesium irradiator at a rate of B166 1.34 cGy min. In tumour-targeted radiation experiments, mice were restrained in a lead shield (Brain Tree Scientific) to minimize exposure to the non-tumour areas.

#### Cell Titer Glo/ Caspase Glo

HCT-116 cells were transfected in a 6 well plate with miR-451a-5p mimic or inhibitor, and the corresponding controls from Life Technologies as previously described. Cells were transferred to a 96 well plate 16 hours post-transfection (1000 cells/well). At 24 hours post-transfection the HCT-116 cells were irradiated with 0, 2, or 5 Gy. Cell Titer-Glo and Caspase 3/7 Glo were analyzed at 48 hours and 96 hours post-irradiation, according to manufacturer’s instructions.

#### RISC trap

HCT116 cells were co-transfected with a plasmid coding for a flag-tagged dominant negative GW418 mutant (Clontech kit #632016) along with a control mimic or miR-451a-5p mimic according to manufacturer instructions. Twenty-four hours later the RNA protein complexes were crosslinked and the RISC complex was immunoprecipitated using an anti-FLAG antibody and RNA was isolated for quantitative real-time PCR of target genes. The fold enrichment was calculated using pre and post IP controls as well as normalization to the control mimic pull-downs.

#### miR in-situ hybridization

Human Colorectal Cancer Tissue arrays (Formalin fixed, # BC05118) were purchased from US Biomax. In-situ hybridization was performed on tissue arrays as described by Pena et al ^29^ using a DIG labeled miR-451a Locked Nucleic Acid (LNA) probe (Exiqon). DIG was detected by an anti-DIG HRP antibody (Roche) and amplified using a TSA-Plus Cy3 system (Perkin Elmer). Tissue arrays were imaged on a Floid Imaging station and scored qualitatively on a 1-4 scale by K.K.

#### Western Blotting

Cell lysates were prepared in RIPA buffer (Pierce 89900) and quantified using a BCA assay (Pierce, #23227) kit. Equivalent amounts of protein were loaded on a 4-15% gradient SDS-polyacrylamide gel (Mini-PROTEAN TGX Precast Gels, BioRad) and transferred onto Nitrocellulose membranes using TransBlot Turbo (BioRad). Membranes were blocked in 5% milk and incubated with antibodies as indicated-CAB39 (Genetex #110628, 1:500 o/n), EMSY (Abcam, #32329 1:300 o/n), Anti-β-actin antibody (Sigma, A5316, 1:10,000 1h RT). Membranes were washed in TBST and incubated with appropriate secondary antibodies from Licor Biosciences (1:15000). Blots were scanned on the Licor Odyssey scanner according to manufacturer’s instructions.

## Author Contributions Statement

R.R, S.R, K.K, C.E-D., C.H designed and performed experiments, analyzed the data. C.L. provided pathology guidelines and samples. C.R.T helped with experimental design and analysis of data. L.V.T and S.A. designed experiments, analyzed data, supervised the study, wrote the manuscript.

## Competing Financial Interests Statement

L.V.T and S.A. are named inventors on a provisional patent application based on some of the findings in this manuscript.

